# Experimental SARS-CoV-2 infection of bank voles - general susceptibility but lack of direct transmission

**DOI:** 10.1101/2020.12.24.424203

**Authors:** Lorenz Ulrich, Anna Michelitsch, Nico Halwe, Kerstin Wernike, Donata Hoffmann, Martin Beer

## Abstract

After experimental inoculation, SARS-CoV-2 infection was proven for bank voles by seroconversion within eight days and detection of viral RNA in nasal tissue for up to 21 days. However, transmission to contact animals was not detected. Therefore, bank voles are unlikely to establish effective SARS-CoV-2 transmission cycles in nature.

**Article Summary Line:** Bank voles show low-level viral replication and seroconversion upon infection with SARS-CoV-2, but lack transmission to contact animals.

---

The severe acute respiratory syndrome coronavirus 2 (SARS-CoV-2) led to a global pandemic in the human population within a few months after its first reporting [1]. Potential wildlife reservoirs of SARS-CoV-2 remain unknown, but susceptibility of various animal species has been described [2, 3]. Among different rodent species, the Syrian hamster (*Mesocricetus auratus*) [4] and the North American deer mouse (*Peromyscus maniculatus*) [5, 6] (both *Cricetidae* species) proved to be highly susceptible. They transmit the virus to co-housed contact animals and are therefore likely to develop effective infection chains in nature. This poses a potential threat, as the development of independent SARS-CoV-2 transmission cycles in nature and a sequentially re-introduction to the human population might be possible [5, 6]. In Europe, bank voles (*Myodes glareolus)* are a wide-spread *Cricetidae* species [7]. Hence, we aimed to characterize the SARS-CoV-2 infection in bank voles and their ability to maintain sustainable infection chains.

Nine bank voles were intranasally inoculated with SARS-CoV-2 strain Muc-IMB-1. Twenty-four hours after experimental inoculation, three contact bank voles were co-housed. Swab samples were taken regularly from all bank voles (Technical Appendix) and one to two animals were euthanized on predefined time-points (4, 8, 12, and 21 days post inoculation (dpi); unfortunately, one bank vole did not survive the initial anesthesia for inoculation).

Neither inoculated nor contact animals showed clinical signs over the course of the study. Seroconversion was detected for all directly inoculated animals sacrificed 8, 12 and 21 dpi, while the animals euthanized 4 dpi and the contact bank voles were all clearly seronegative for SARS-COV-2 antibodies in an already validated, indirect, multi-species ELISA based on the receptor-binding domain (RBD) [8].

All directly inoculated bank voles tested RT-qPCR-positive for SARS-CoV-2 in the oral and rhinarium swabs 2 dpi. At 4 dpi, five of these eight bank voles were positive in oral swabs. Two of them were additionally positive in the rhinarium swabs. On both mentioned sampling days, rectal swabs of two animals each tested RT-qPCR positive for SARS-CoV-2. Groupwise-collected fecal samples also tested RT-qPCR positive on 2 and 4 dpi. All swabs collected 8, 12 and 16 dpi from directly inoculated animals and every swab from the co-housed contact animals tested RT-qPCR negative. For details see Figure 1 and Table 1.

**Figure 1.**
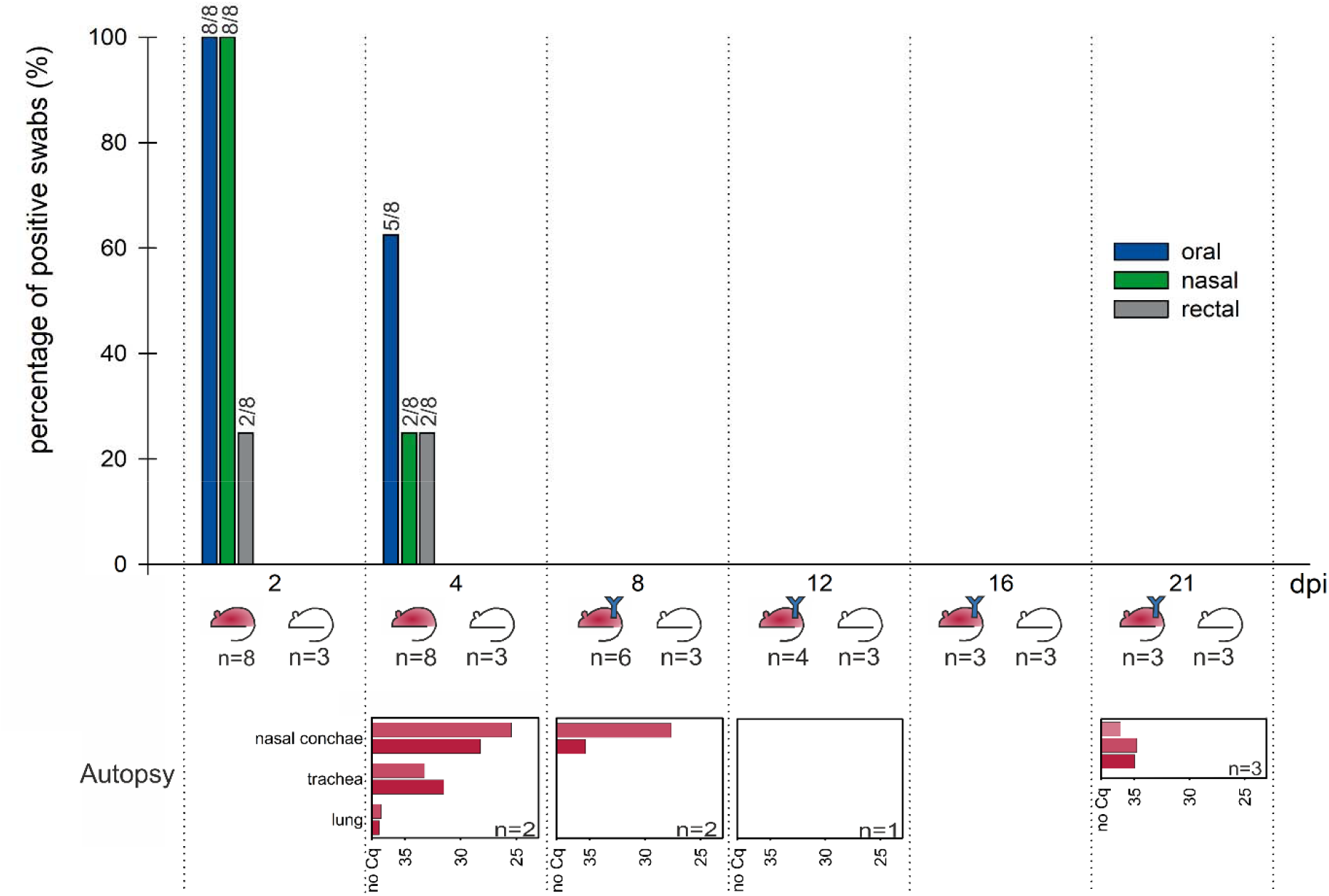
Percentage of RT-qPCR positive swabs on all sampling time-points. The “red mouse symbol” symbolizes inoculated bank voles, while the “white mouse symbol” represent co-housed contact bank voles. Blue “Y” symbols stand for detected antibodies against SARS-CoV-2 in the respective bank vole group. RT-qPCR results for the sampled organs of the euthanized, inoculated bank voles are given below the main chart for each time-point. n: number of bank voles; dpi: days post inoculation; Cq: quantification cycle

**Table 1.**
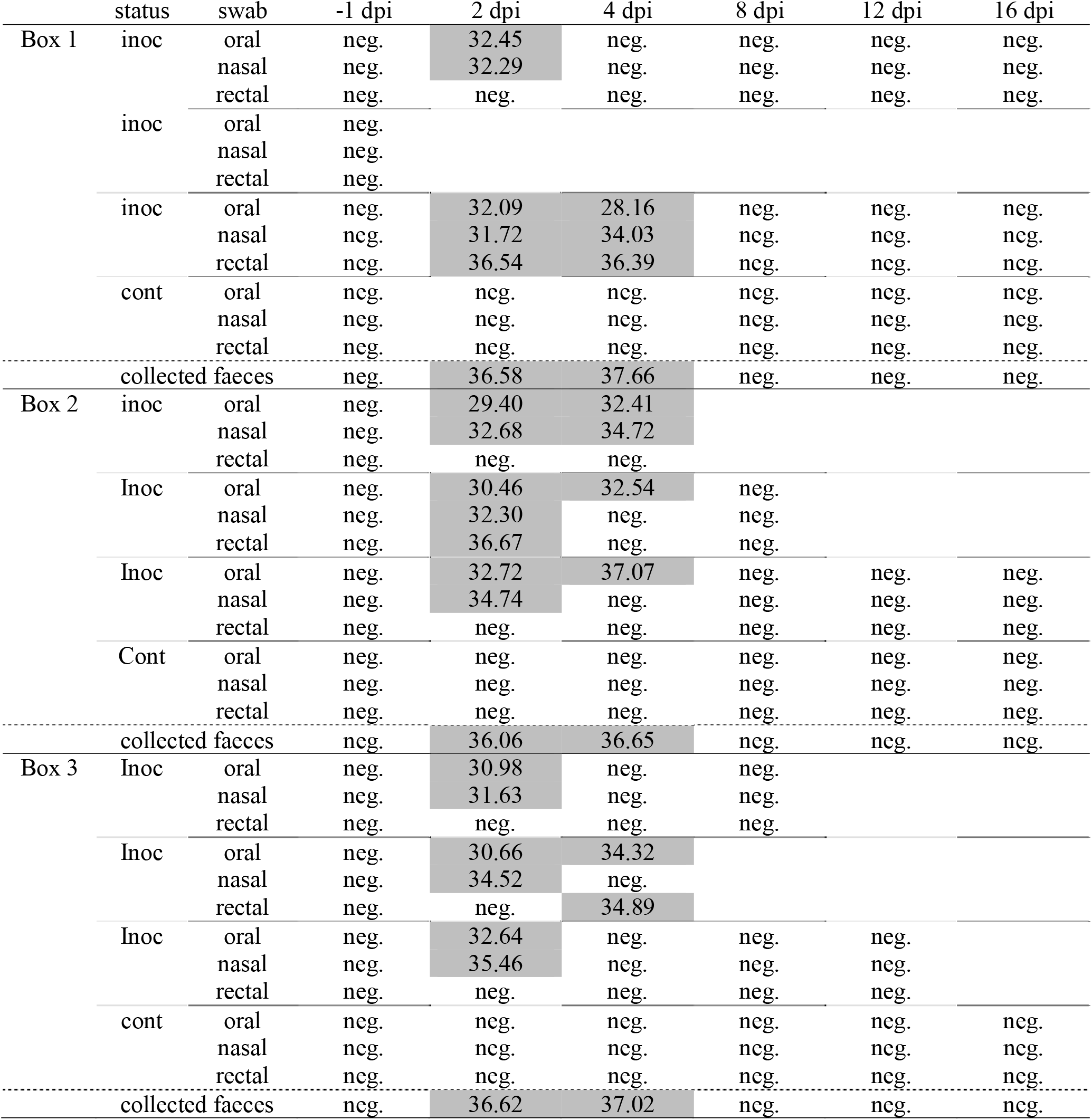
RT-qPCR results of the swap sampling of all inoculated (inoc) and contact (con) bank voles. Results are given in quantification cycle values (Cq). dpi: days post inoculation

Two animals were sacrificed on 4 dpi. The nasal conchae, trachea, lung and olfactory bulb tested RT-qPCR positive for SARS-CoV-2 RNA (quantification cycle (Cq) values 25.45-37.15). One animal showed viral genome in cerebrum and cerebellum samples, while the other one was positive in the spleen sample. At 8 dpi another two animals were sacrificed. Both exhibited viral RNA only within the nasal conchae. The animal sacrificed 12 dpi was negative in all collected tissue samples. The three inoculated animals euthanized 21 dpi tested RT-qPCR positive in the nasal conchae (Cq values 34.78, 34.97, 36.25), while the three contact animals euthanized at the same time-point tested all negative in the nasal conchae.

Re-isolation of viable virus from tissue materials in cell culture (Vero E6) was successful for one nasal conchae sample taken at 4 dpi. However, isolation from samples with Cq >28 failed, which is in line to findings of other groups [3, 9].

Overall, bank voles proved to be susceptible for an infection with SARS-CoV-2, but do not transmit the virus to co-housed direct contact animals. The presented results suggest a tissue tropism for SARS-CoV-2 replication in bank voles to the upper respiratory tract, as likewise seen for other species, e.g. ferrets, fruit bats, and raccoon dogs [3, 9]. The persistence of viral genome for at least three weeks in nasal tissue of directly inoculated animals was unexpected, especially since the last RT-qPCR positive swab was retrieved 4 dpi from the respective bank voles (Table 1). This is most likely due to the suspected clustering of SARS-CoV-2 infection foci in narrow areas of the upper respiratory tract [10]. Since virus isolation from these 21 dpi samples was not successful, the persistence of SARS-CoV-2 is unlikely to lead to the same shedding of infectious virus as it was shown before for deer mice [5, 6]. Additionally, deer mice seem to shed virus also through the rectum. However, in bank voles SARS-CoV-2 genome could not be detected in the intestines. Even though rectal swabs and fecal samples were RT-qPCR positive, the detected Cq were high, indicating low viral RNA levels. Therefore, the detected viral RNA likely represents residues, which might have resulted from extensive grooming behavior and therefore do not correspond with actual virus shedding from the rectum or feces.

This study proves a general susceptibility of bank voles towards SARS-CoV-2 infection. However, bank voles did not transmit SARS-CoV-2 to contact animals in the presented study. This makes them unlikely to maintain sustainable infection chains in nature. Therefore, the risk of bank voles becoming a reservoir for SARS-CoV-2 in nature, e.g. after contact to infected cats, is unlikely.

## Acknowledgments

We thank Mareen Lange, Anke Eggert, Bianka Hillmann, Frank Klipp, Doreen Fiedler and Harald Manthei for their excellent assistance in the lab and dedicated animal care. This research was supported by intramural funding of the German Federal Ministry of Food and Agriculture provided to the Friedrich-Loeffler-Institut and partial funding from the European Union Horizon 2020 project (“Versatile Emerging infectious disease Observatory”, grant no. 874735).

## Ethical Statement

The experimental protocol was assessed and approved by the ethics committee of the State Office of Agriculture, Food Safety, and Fisheries in Mecklenburg-Western Pomerania (permission number MV/TSD/7221.3-2-010/18).

## Technical Appendix

## Animals and housing conditions

Eight female and four male bank voles were obtained from an in-house breeding colony at the Friedrich-Loeffler-Institut, Insel Riems, Germany. The age ranged from 7 to 9 weeks. Prior to infection, a negative serological status towards SARS-CoV-2 of the breeding colony was determined by an indirect RBD-ELISA (1). Further, all animals used in the trial were tested RT-qPCR negative for SARS-CoV-2 one day prior to infection by rhinarium, oral and rectal swabs. For the duration of the study the animals were kept in individually ventilated cages (IVC) with a light regime of 12 hours illumination and 12 hours darkness. Drinking water and a rodent diet were provided ad libitum. All handling procedures were performed under BSL-3 conditions.

## Study design

Nine bank voles were inoculated with 1×10^5^ tissue culture infection dose 50 (TCID_50_) of the SARS-CoV-2 strain “2019_nCoV Muc-IMB-1” (GISAID ID_EPI_ISL_406862, designation “hCoV-19/Germany/BavPat1/2020”) by administering 70 μl virus suspension to the nostrils and rhinarium. Inoculation took place under a short-term isoflurane based inhalation anesthesia. Three inoculated bank voles were housed together in one IVC each. Twenty-four hours after inoculation another three - one per IVC - naïve in-contact bank voles were co-housed with the directly inoculated animals. A physical examination following a defined clinical score regarding general behavior, respiration, eyes and neurologic symptoms was performed daily and body weight changes were monitored regularly (0, 2, 3, 4, 6, 7, 8, 9, 10, 12, 14, 16, 21 days post infection [dpi]). Oral, rhinarium and cloacal swabs were taken from each animal at 2, 4, 8, 12, 16 dpi. Further, a fecal sample was taken from each IVC at the aforementioned sampling points.

Two bank voles each were sacrificed at 4 and 8 dpi and another one at 12 dpi. At autopsy, a serum sample was collected and the nasal conchae, trachea, lung, heart, olfactory bulb, forebrain, cerebellum, liver, spleen, kidney, small and large intestine were sampled. The remaining animals were euthanized at 21 dpi and serum samples were collected as well as a sample of the nasal conchae.

## Antibody detection

Serum samples were tested by the aforementioned RBD-ELISA (1). Absorbance values larger than 0.3 are considered antibody positive, those lower than 0.2 antibody negative, and in between as questionable. The results are presented in Supplementary Table 1.

## RNA extraction and RT-qPCR

Before sampling, swabs (nerbeplus GmbH&Co KG, Germany and Copan Italia S.p.A., Italy) were dampened with Hank’s 692 balanced salts (HBS) and Earle’s balanced salts (EBS) in minimum essential medium (MEM) After sampling, the swabs were resuspended in 1 ml HBS and EBS MEM with the addition of penicillin and streptomycin. Fecal samples were directly collected in 1 ml of HBS and EBS MEM with the addition of penicillin and streptomycin. Organ samples were transferred in 1 ml of HBS and EBS MEM with an added steel bead and homogenized at 30,000 Hz for two minutes with the TissueLyserII (Qiagen, Germany). Nucleic acid was extracted from 100 μl of the supernatant of all samples with the NucleoMag Vet kit (Macherey-Nagel, Germany). Extracted viral RNA levels were determined by the already validated RT-qPCR “nCoV_IP4”, targeting the viral RNA-dependent RNA polymerase (2). A quantification cycle (Cq) value of 38 was used as a cut-off value. The results are presented in Supplementary Table 1.

## Virus isolation

Virus re-isolation in cell culture was attempted on a Vero E6 cell line (L0929, collection of cell lines in veterinary medicine, Insel Riems, Germany) using HBS and EBS MEM with the addition of penicillin and streptomycin. Viral replication was determined by cytopathic effect (cpe) within 72 hours after inoculation. Cultures with no visible cpe in the first passage were passaged once.

---

**Supplementary Table 1.**
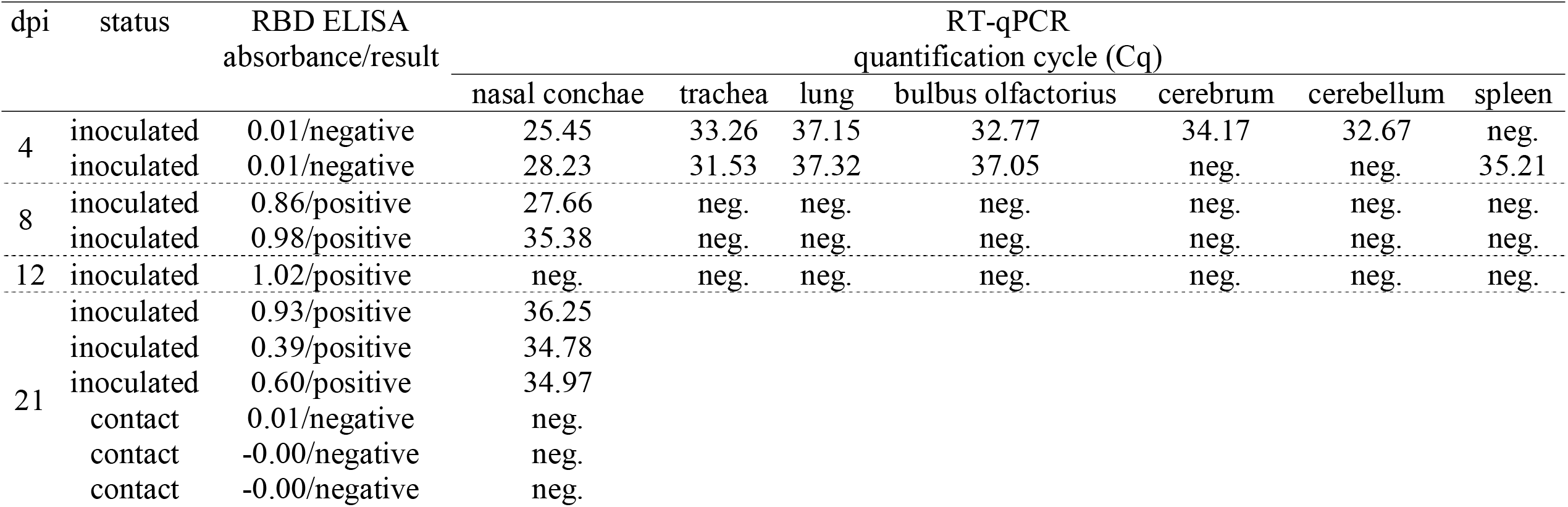
RT-qPCR results of the organ samples of all inoculated and contact bank voles as well as results from the indirect, multispecies ELISA. RT-qPCR results are given in quantification cycle values (Cq). dpi: days post inoculation

